# Lysine specific demethylase 1 (LSD1) regulates host alpha-ketoglutarate levels to modulate lipid peroxidation during *Mycobacterium tuberculosis* infection

**DOI:** 10.64898/2026.04.20.719577

**Authors:** Awantika Shah, Aakash Chandramouli, Atheena Abhayakumar, Raju S Rajmani, Siddhesh S Kamat, Kithiganahalli Narayanaswamy Balaji

## Abstract

*Mycobacterium tuberculosis* (Mtb) subverts host immune responses via modulation of host epigenome and metabolism. In this study, we underscore a role for the epigenetic modifier, Lysine Specific Demethylase 1 (LSD1), in regulating macrophage metabolism to support mycobacterial pathogenesis. In *ex vivo* and *in vivo* infection models, LSD1 inhibition reduced mycobacterial CFU alleviating lung pathology. Metabolomic analysis of Mtb infected, LSD1 deficient macrophages revealed increased levels of alpha-ketoglutarate (AKG), a crucial TCA cycle metabolite via regulating genes implicated in glutamine breakdown. Moreover, exogenous addition of AKG resulted in reduced oxidative stress and attenuated lipid peroxidation (LPO) with a consequent decrease in Mtb survival. Blocking glutamine breakdown in LSD1 deficient macrophages failed to reduce LPO and promoted Mtb intracellular survival, highlighting the role of LSD1-AKG axis in this immunomodulation. Dietary supplementation of AKG to Mtb infected mice improved lung pathology, limited Mtb dissemination and reduced the levels of oxidative Malondialdehyde adducts. Therefore, we highlight a host protective role of AKG during Mtb pathogenesis through suppression of lipid peroxidation and uncover an epigenetic-metabolic axis exploited by Mtb, thereby positing dietary supplementation of AKG as a potential therapeutic strategy against Tuberculosis.

## INTRODUCTION

Modulation of host epigenome as a pathogen strategy to manipulate host responses has been characterized in various infections^1,2^. *Mycobacterium tuberculosis* (Mtb) is also known to exploit host epigenetic machinery towards its benefit via histone modifications, DNA methylation and miRNA mediated transcriptional changes^3,4^. Histone demethylases are key players in innate and adaptive immunity with roles in controlling maturation, differentiation and functions of immune cells^5^. Downregulation of histone demethylase KDM6B leading to H3K27 hypermethylation has been associated with active pulmonary TB disease^6^. Foam cell formation induced by histone demethylase JMJD3 has been shown to aid mycobacterial survival^7^. Another histone demethylase, LSD1/KDM1A, (Lysine Specific Demethylase 1) has emerged as a key regulator of macrophage inflammatory responses^8,9^, lipid metabolism^10,11^ as well as cellular metabolism^12,13^. With well documented roles in governing cancer progression and regulating viral and fungal infections^14–18^, LSD1 posits as a potential therapeutic target with various LSD1 inhibitors undergoing clinical trials^19–21^. However, a role for LSD1 in governing Mtb pathogenesis is not yet explored.

In this study, we show that LSD1 regulates levels of host alpha-ketoglutarate (AKG), a key metabolite of TCA cycle, to support Mtb infection. AKG can be derived from two sources, either during TCA cycle by oxidative decarboxylation of isocitrate or via breakdown of the amino acid, glutamine^22^. AKG, generally associated with promoting anti-inflammatory responses in macrophages, is known to alleviate various disease pathologies^23^. Studies have implicated an antioxidative function of AKG during cardiovascular and age-related diseases, cancers as well as neurological disorders^24^. During Mtb infection, excessive oxidative stress driven lipid peroxidation (LPO) has been implicated in supporting dissemination and pathogenesis via mediating ferroptotic cell death pathway ^25–28^. Given a role of AKG in resolving inflammation via antioxidant properties and inhibiting ferroptosis^29,30^, we hypothesized it might play a role in counteracting Mtb-induced LPO and investigated the role of the epigenetic modifier, LSD1, in this immunomodulation.

Here, we report that Mtb infection leads to increased expression of LSD1 which was found to be indispensable for its survival within the host milieu. Knocking down LSD1 in macrophages led to increased levels of AKG and transcriptional regulation of genes implicated in glutamine utilization. Further, exogenous addition of AKG led to decreased intracellular survival of Mtb via inhibiting Mtb-induced cellular reactive oxygen species (ROS) levels and LPO. LSD1 inhibition mediated regulation of Mtb pathogenesis was dependent on AKG production as upon blocking glutamine breakdown, there was increased LPO and enhanced mycobacterial survival in LSD1 deficient macrophages. Finally, dietary supplementation of AKG to Mtb-infected mice alleviated TB pathology with a concomitant decrease in lung bacillary burden. Levels of Malonaldehyde (MDA) adducts, by-product of lipid peroxidation, were reduced in lungs of mice given AKG with a subsequent decrease in bacterial dissemination. Altogether, our study uncovers a role of LSD1 in modulating macrophage metabolism during Mtb infection and positions AKG supplementation as a potential therapy against Tuberculosis (TB).

### Histone demethylase LSD1 is upregulated upon Mtb infection

A role for LSD1 in modulating host cellular pathways and metabolism during cancers and infections is well reported^17,31–33^. To understand its implications during Mtb pathogenesis, we first assessed for its regulation during infection and observed increased transcript levels of LSD1 (*Kdm1a*) in Mtb H37Rv infected macrophages 6h and 12h post infection **(Figure 1A)**. In corroboration, protein levels of LSD1 were upregulated in Mtb H37Rv infected macrophages **(Figure 1B-E)**. In an *in vivo* mice model of TB infection, an increase in the expression of LSD1 in lungs of Mtb infected mice at transcript **(Figure 1G)** as well as protein levels **(Figure 1H)** was observed. Together, our results suggest that upon infection, Mtb induces host epigenetic modifier, LSD1 during both, *ex vivo* and *in vivo* infections.

**Figure 1:**
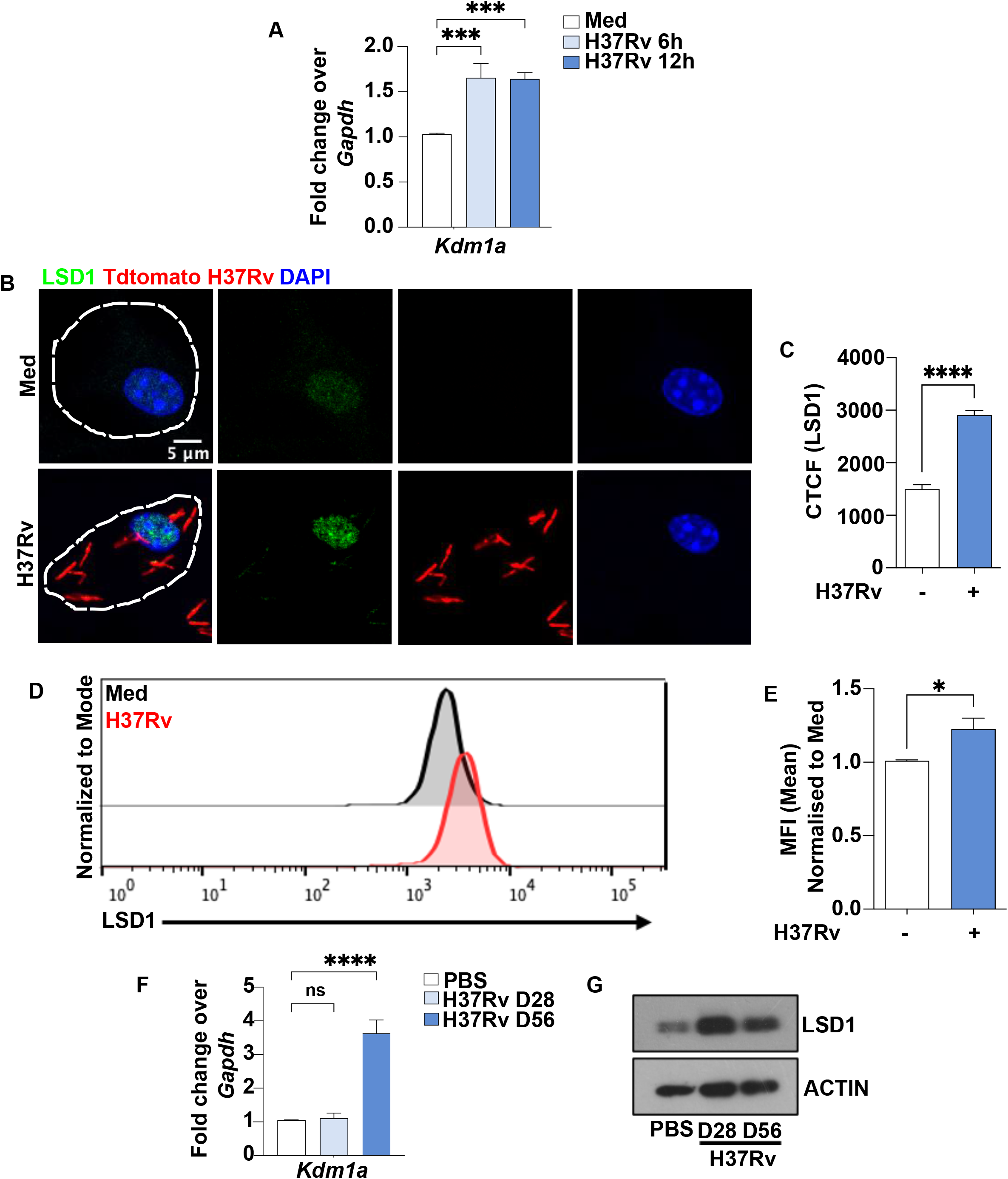
Histone modifier, LSD1, is upregulated during *Mycobacterium tuberculosis* infection. Mtb H37Rv infection was given to macrophages for the mentioned timepoints and expression of LSD1 was assessed at **(A)** transcript levels and protein levels by **(B-C)** Immunofluorescence and **(D-E)** Flow cytometry. Mice infected with H37Rv via aerosol route (100 CFU) were assessed for LSD1 levels in lungs post 28 and 56 days at **(F)** transcript and protein levels by **(G)** Immunoblotting. CTCF-Corrected Total Cell Fluorescence, ns, non-significant, *, P< 0.05 ***, p<0.0005 ****, p<0.0001 Student’s t-test: A, C, E and F. N=3.

### Mtb induced LSD1 aids mycobacterial pathogenesis

Next, we sought to investigate if Mtb mediated increase in host LSD1 levels have any role in governing mycobacterial pathogenesis. To this end, siRNA-mediated knockdown of LSD1 was carried out in mice macrophages **(Supplemental Figure 1A)** and assessed for Mtb survival. A decrease in intracellular Mtb CFU was observed in macrophages wherein LSD1 was knocked down **(Figure 2A)** as well as upon pharmacological inhibition of LSD1 using a specific monoamine oxidase (MAO) inhibitor **(Supplemental Figure 1B)**. To evaluate the same during *in vivo* TB infection, mice infected with H37Rv via aerosol route were intraperitoneally given LSD1 inhibitor treatment **(Figure 2B)**. LSD1 inhibition resulted in reduced Mtb burden in mice lungs as compared to vehicle control **(Figure 2C)**. Further, when administered in combination with Isoniazid, a frontline TB drug, LSD1 inhibition resulted in enhanced mycobacterial clearance from mice lungs, highlighting a pro-pathogen role of LSD1 during Tuberculosis. This synergistic effect was also evident in lung pathology, which was improved upon combined LSD1 inhibition and isoniazid treatment, as assessed by gross lung pathology **(Figure 2D)**, granuloma scoring **(Figure 2E)** and H/E staining **(Figure 2F)**. Altogether, our findings indicate that induction of LSD1 by Mtb could be a strategy to support its pathogenesis.

**Figure 2:**
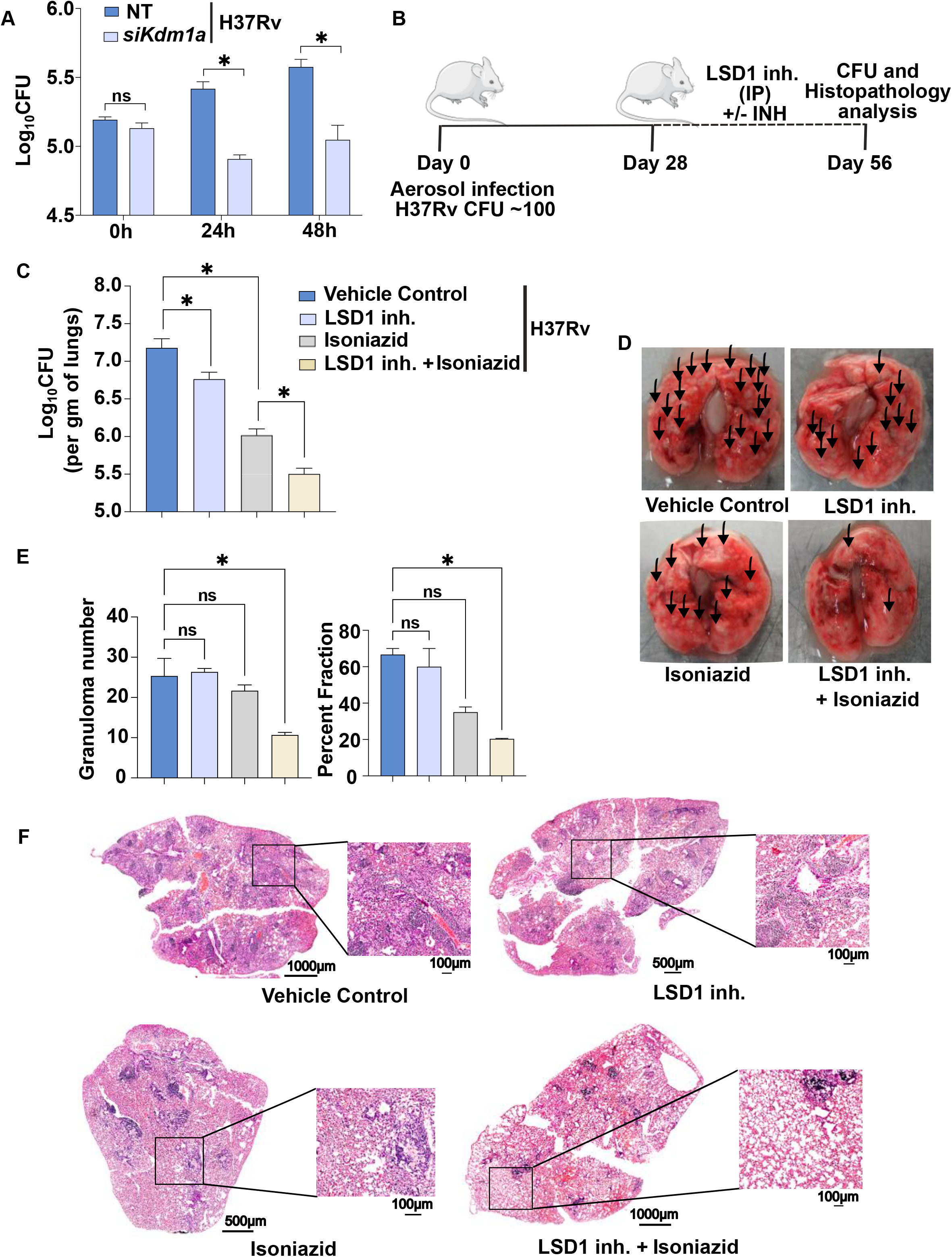
LSD1 aids mycobacterial pathogenesis. **(A)** In vitro CFU was assessed in mouse macrophages post transfection with siRNAs against *Kdm1a* for mentioned time points of H37Rv infection (MOI 1:5). **(B)** Mice given aerosol infection with 100 CFU of H37Rv for 28 days followed by for intraperitoneal LSD1 inhibitor (50mg/kg) or Isoniazid (INH) (10mg/kg) therapeutic treatment. At 56 day, Lungs of mice from the mentioned groups were processed for **(C)** CFU **(D)** gross pathology, H and E staining - **(E)** Granuloma number (left), Percent fraction of lungs covered by granuloma (right), **(F)** representative image. Inh.-inhibitor, IP-intraperitoneal. ns, non-significant *, p<0.05 Student’s t-test : A. *, p<0.05 One-Way ANOVA: C and E. N=3.

### LSD1 knockdown leads to increase in levels of Alpha-ketoglutarate during Mtb infection

Mtb-induced LSD1 could be involved in modulation of various host cellular processes governing Mtb pathogenesis and is known to facilitate tumour adaptation to hostile microenvironments via metabolic reprogramming and regulating gene expression to facilitate cancer progression^12,34^. Furthermore, metabolically distinct macrophage populations during Mtb infection exhibit differential capacities to control Mtb burden^35^.

To understand if LSD1 alters host metabolism to support mycobacterial growth, we performed a focused metabolomics analysis upon LSD1 KD during Mtb infection and first assessed for TCA cycle metabolites. Metabolic extraction and metabolomics analysis was carried out in Mtb infected mice macrophages wherein LSD1 was knocked down as per established protocol^36,37^. Although, levels of several TCA cycle intermediates were altered upon Mtb infection, most of the metabolites were not regulated by LSD1 indicating that LSD1 might not be modulating TCA during Mtb infection **(Figure 3A)**. Interestingly, cellular concentration of 2-ketoglutarate (2-KG) (or α-ketoglutarate, AKG), an important TCA metabolite^38^, was upregulated upon LSD1 KD in Mtb infected macrophages.

**Figure 3:**
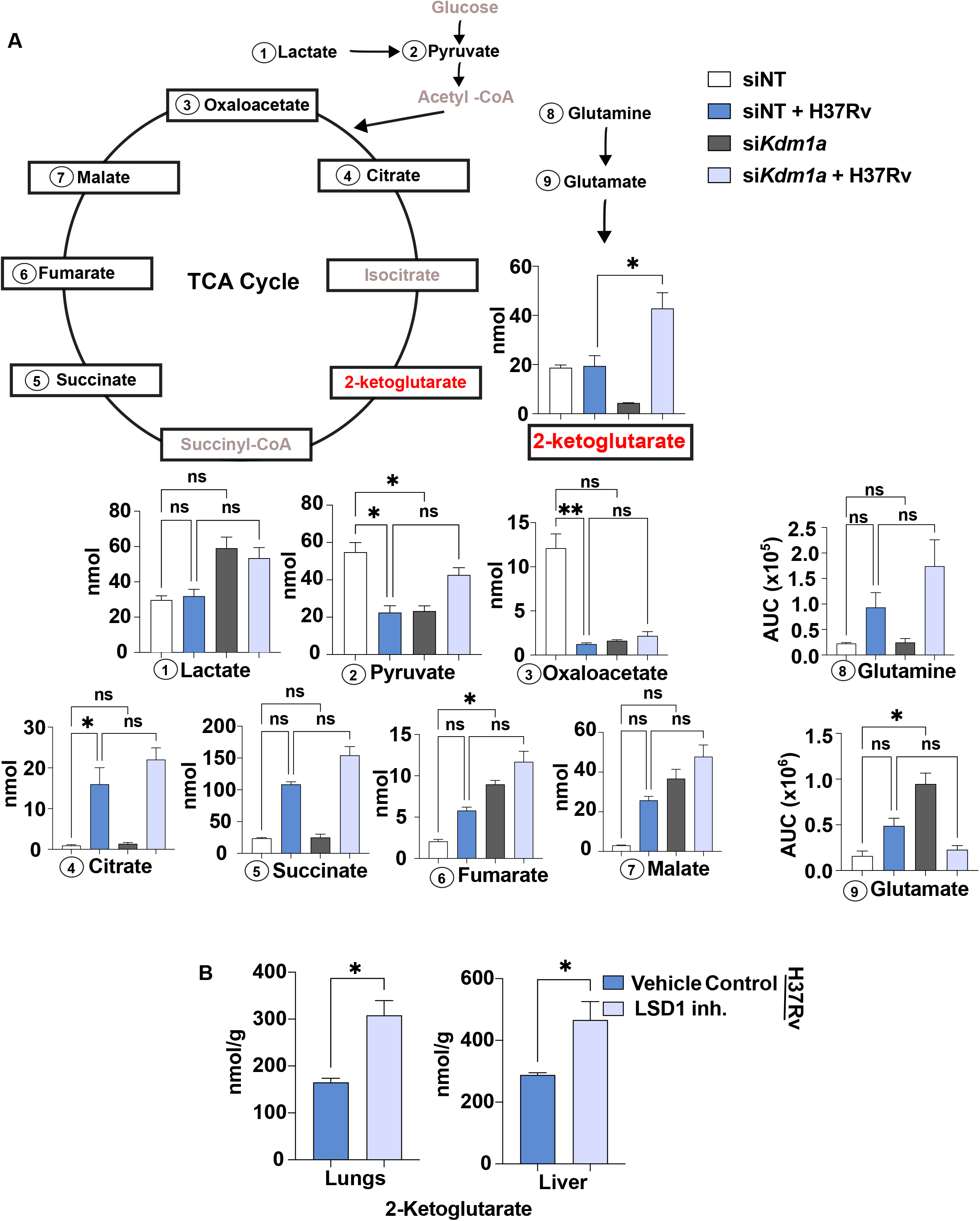
Mtb-induced LSD1 modulates host AKG levels. **(A)** Transient transfection was carried out in mice macrophages with non-targeting (NT) or *Kdm1a* siRNA. 24h infection with H37Rv was given and levels of various metabolites of TCA cycle were assessed via targeted mass spectrometry. **(B)** AKG levels of Mtb-infected and LSD1 inhibitor (50mg/kg) mice was assessed in mice Lungs (left) and Liver (right) post 56 days of infection and treatment regime. ns, non-significant *, p<0.05 **, p<0.005 One-Way ANOVA: A. *, p<0.05 Student’s t-test: B. N=3.

To gain an *in vivo* perspective, lung and liver of Mtb-infected and LSD1 inhibitor-treated mice were subjected to a similar targeted metabolomics analysis to assess whether regulation of AKG levels observed in *ex-vivo* infection scenario also holds true during TB pathogenesis. Indeed, we observe upregulation of AKG upon LSD1 inhibition in lungs and liver of mice **(Figure 3B)**, corroborating with improved lung pathology upon LSD1 inhibitor treatment. Metabolites have also been reported to regulate immune functions in addition to their roles in cellular energy metabolism^39^. This aligns with previous studies highlighting a role for AKG in mitigating lung inflammation and reducing fatty liver disease progression^40–43^. Overall, our data suggest that during Mtb infection, LSD1 mediated regulation of AKG levels might have implications in facilitating TB pathogenesis.

### LSD1 KD enhances glutamine breakdown to regulate AKG levels during Mtb infection

Synthesis of AKG occurs via oxidative carboxylation of isocitrate via isocitrate dehydrogenase during TCA cycle or via breakdown of glutamine (Glutaminolysis), with the latter being reported as the main source of AKG in macrophages during infection^23^. To probe into LSD1 mediated regulation of AKG, we wanted to assess for modulation, if any, of these pathways during Mtb infection. We performed ^13^C_6_ glucose tracing experiment to assess metabolic flux post Mtb infection in control and LSD1 siRNA treated macrophages. A comparative analysis between ^13^C_6_ labelled (Glucose derived) and ^12^C_6_ metabolite pools post normalization to uninfected condition would give us an idea how LSD1 might be affecting substrate utilization during Mtb infection. We assessed for levels of AKG derived from TCA cycle (Glucose-^13^C_6_) or via glutaminolysis (Glutamine-^12^C_6_) across mentioned time points. Interestingly, we observe an increase in both ^12^C_6_ and ^13^C_6_ derived AKG in LSD1 KD macrophages during Mtb infection **(Figure 4A)**.

**Figure 4:**
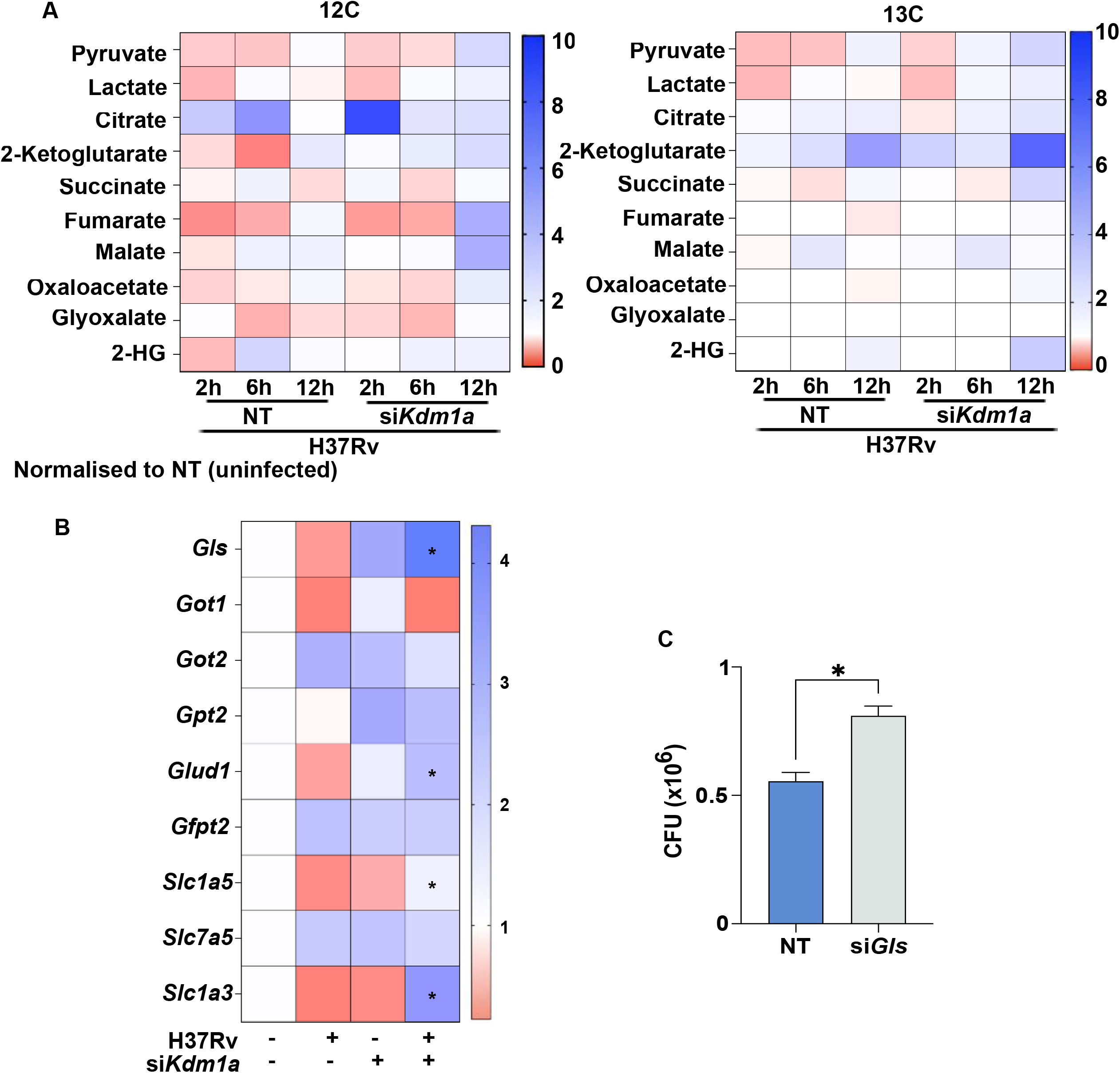
Knocking down LSD1 results in increased glutamine breakdown upon Mtb infection. (A) Metabolic flux analysis carried out using ^13^C_6_ glucose tracer experiments for 2,6 and 12h in mice macrophages transiently transfected with either non-targeting (NT) or siRNA against *Kdm1a* infected with H37Rv for 24h. (B) Mice macrophages transiently transfected with either NT siRNA or siRNA against *Kdm1a* were infected with H37Rv for 24h and transcript levels of mentioned genes were assessed. (C) Mouse peritoneal macrophages transiently transfected with siRNA against *Gls1* and CFU was assessed 48h post H37Rv infection (MOI 1:5). *p< 0.05 One Way ANOVA for B; *, p< 0.05 Student’s t-test for C.

Furthermore, we observed LSD1 dependent transcriptional regulation of various genes of the glutaminolysis pathway **(Figure 4B)**. Increased expression of *Glutaminase (Gls)* and *Glutamate dehydrogenase (Glud1)*, implicated in Glutamine to AKG conversion, and glutamine transporters *(Slc1a5, Slc1a3)* indicates glutamine breakdown in LSD1 deficient macrophages upon Mtb infection. Enhanced intracellular survival of Mtb upon inhibiting glutaminolysis and consequent reduction in AKG production, via knocking down *Gls* in macrophages **(Supplemental Figure 3A, Figure 4C)**, suggests a role for glutamine derived AKG in restricting mycobacterial survival.

Taken together, our results indicate that LSD1 acts as a metabolic regulator that supports Mtb pathogenesis as LSD1 KD in macrophages led to elevated AKG levels, likely through enhanced glutamine breakdown and inhibition of this process resulted in increased mycobacterial survival.

### AKG supplementation reduces mycobacterial survival via suppressing host lipid peroxidation

Given the current line of investigation, it becomes crucial to assess whether exogenous addition of AKG could be beneficial to host during Mtb infection. To this end, we carried out an *ex vivo* Mtb CFU enumeration and macrophage cell death assay upon AKG supplementation. Intracellular survival of Mtb in murine macrophages was decreased upon AKG supplementation **(Figure 5A)** and Mtb induced macrophage cell death was reduced upon AKG supplementation **(Figure 5B)**, hinting towards an overall host protective role of AKG during Mtb infection.

**Figure 5:**
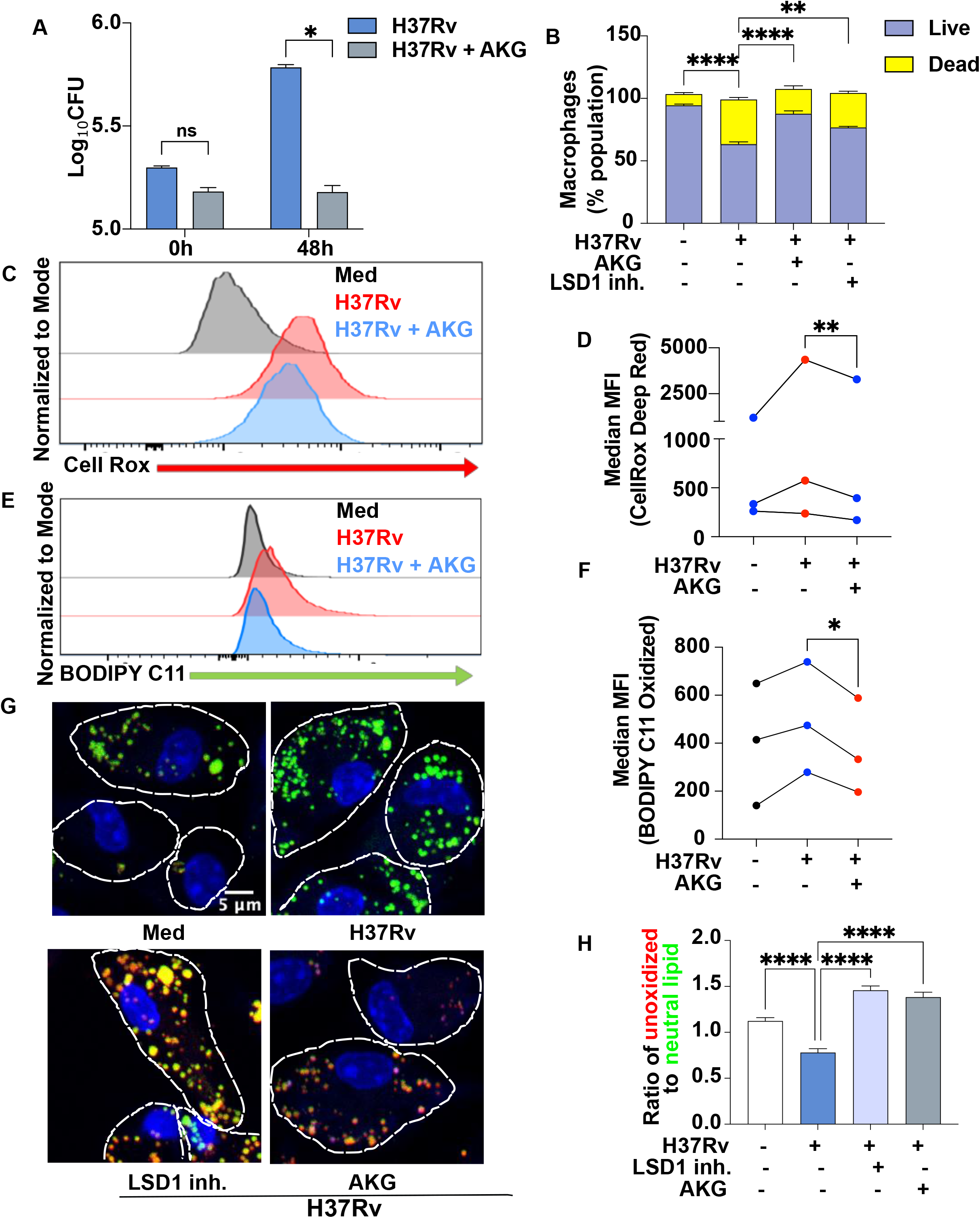
Alpha-KG reduces Mtb survival via modulating host lipid peroxidation. **(A)** *In vitro* CFU was assessed in mice macrophages infected with H37Rv (MOI 1:5) with and without 2mM exogenous alpha-ketoglutarate (AKG) post 48h of infection. **(B)** Mice macrophages infected with H37Rv and LSD1 inhibitor treatment or AKG supplementation were assessed for macrophage cell death via PI staining. **(C-F)** Mice macrophages post H37Rv infection for 24h in presence and absence of AKG supplementation were analysed for **(C**,**D)** oxidative stress via CellRox DeepRed stain and **(E**,**F)** lipid peroxidation by BODIPY C11 staining via flow cytometry. **(G-H)** Lipid peroxidation was assessed using BODIPY 665 and BODIPY 493/503 dye in 24h H37Rv infected mice macrophages with and without LSD1 inhibitor treatment or AKG supplementation **(G)** Representative image, **(H)** quantification. ns, non-significant *, p< 0.05 **, p<0.005 Student’s t-test: A, D and F. **, p< 0.005 ****, p< 0.0001 One-Way ANOVA: B, H. N=3.

During infections, AKG is generally implicated in reducing host stress and resolving inflammation via its antioxidant activity^44–46^. In addition to being a direct quencher of hydrogen peroxide, AKG promotes expression of anti-oxidant genes, promote synthesis of Glutathione and increases GPX4 levels, a key antioxidant enzyme implicated in reducing lipid peroxidation^24^. Indeed, we found a reduction in macrophage Reactive Oxygen Species (ROS) levels upon AKG supplementation during Mtb infection as measured by CellRox staining **(Figure 5C, D)**. Excessive oxidative stress in the cells lead to lipid peroxidation (LPO) which is implicated in aiding Mtb dissemination and pathogenesis^26^. Mtb-induced lipid peroxidation was decreased in macrophages upon AKG supplementation as evident by BODIPY C11 staining (indicator of lipid oxidation) **(Figure 5E, F)**. AKG supplementation also resulted in increased transcription of GPX4, indicating AKG regulates GPX4 to suppress LPO during Mtb infection (**Supplemental Figure 2A)**. This was concomitant with reduction in Malondialdehyde (MDA) levels, oxidative adducts formed as a by-product of lipid peroxidation, upon AKG supplemented in Mtb-infected macrophages (**Supplemental Figure 2B)**. Finally, upon quantifying oxidized vs reduced lipid levels in macrophages upon Mtb infection, we found an increase in the abundance of reduced lipids upon AKG supplementation as well as upon LSD1 inhibition in Mtb infected macrophages **(Figure 5G, H)**.

In summation, our data suggests AKG might aid mycobacterial clearance via regulating GPX4 levels leading to reduced lipid peroxidation that has been implicated in promoting Mtb dissemination and survival^25–28^. Consistent with this, AKG has been reported to reduce MDA accumulation to enhance cellular antioxidant capacity^24^ as well as inhibit ferroptosis associated LPO to alleviate osteoarthritis^29,30^. Therefore, AKG might be potentiating anti-TB responses via reducing Mtb driven host lipid peroxidation and oxidative stress.

### LSD1-AKG axis is implicated in regulating host lipid peroxidation and Mtb survival

Our data suggests that LSD1 inhibition during Mtb infection increases host AKG levels, decreases LPO and restricts mycobacterial survival. In line with this, exogenous supplementation of AKG also results in similar reduction of LPO and Mtb CFU. To determine if this effect is a consequence of LSD1 mediated regulation of AKG, we knocked down *Gls1*, to block glutamine breakdown and consequent AKG production, along with LSD1 knockdown. Specifically, we carried out intracellular Mtb survival assay upon siRNA mediated knockdown of either LSD1 (*Kdm1a*) or Glutaminase (*Gls1*) or both in Mtb infected macrophages.

Notably, the reduction in Mtb CFU observed upon LSD1 knockdown was abrogated upon double knockdown of LSD1 and *Gls1* suggesting that protective effect observed upon LSD1 inhibition is dependent on glutamine derived AKG **(Figure 6A)**. We further assessed for the status of reduced vs oxidized lipids during Mtb infection and observed that the increase in reduced lipid content upon LSD1 knockdown was reversed upon *Gls* knockdown, both alone and in combination with LSD1 knockdown **(Figure 6B-C)**. This suggests that LSD1 mediated regulation of host lipid peroxidation during Mtb infection requires *Gls* activity. Therefore, Mtb-induced LSD1 aids mycobacterial survival by regulating lipid peroxidation via metabolic regulation of AKG.

**Figure 6.**
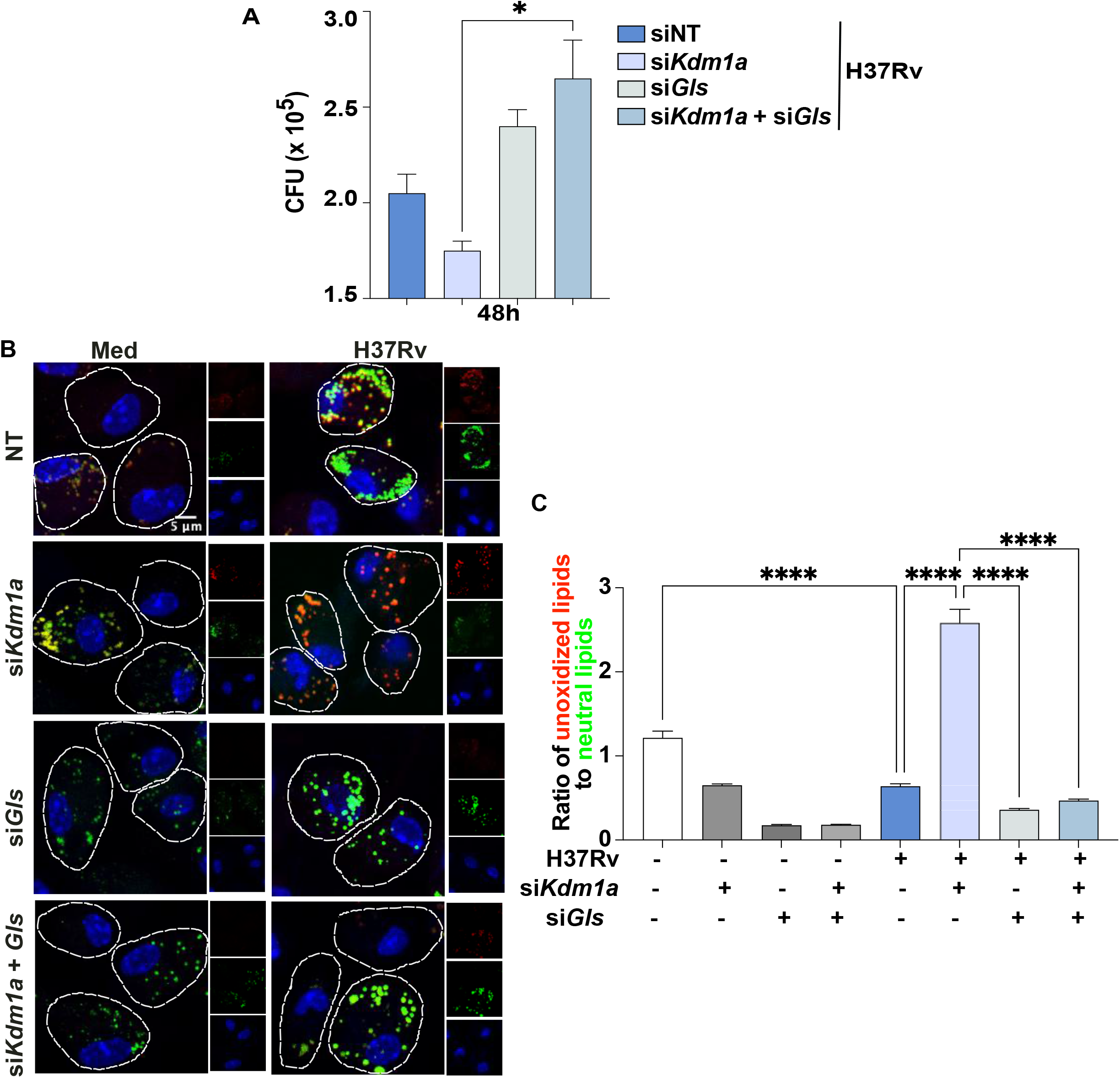
LSD1-AKG axis is involved in regulating host lipid peroxidation during Mtb infection. (A-C) Mice macrophages transfected with either non-targeting siRNA or siRNA against *Kdm1a* or *Gls1* or both were infected with H37Rv and assessed for (A) intracellular Mtb survival at 48h post infection (MOI 1:5), (B-C) lipid peroxidation via BODIPY 665 and BODIPY 493/503 staining. (B) Representative image (C) quantification. *,p<0.05 ****, p< 0.0001 One-Way ANOVA for A and C.

### Dietary supplementation of AKG alleviates TB pathogenesis

Given our findings that AKG was able to suppress Mtb-induced lipid peroxidation which is associated with increased Mtb pathogenesis and dissemination^47^, we wanted to explore the therapeutic potential of AKG during TB pathogenesis. Dietary supplementation of AKG has shown promising results against cancer^48^, inflammatory diseases^49–51^, fatty liver diseases^41^, diabetes^52^ as well as in alleviating lung injuries^43^, SARS COV2 infection^42^ and has been linked with increasing lifespan^53^ and overall health^54,55^. To this end, 4 weeks post aerosol infection with Mtb H37Rv, mice were given AKG supplementation (1% in drinking water) for 4 weeks **(Figure 7A)**. Upon gross lung examination, we observe a decrease in the number of lesions in lungs of mice given AKG supplementation as compared to untreated infected mice **(Figure 7B)**. In line with this, there was a reduction in the lung Mtb burden upon AKG supplementation **(Figure 7C)**. Furthermore, upon assessing for effect of AKG on Mtb dissemination, we found a significant decrease in Mtb CFU in spleen **(Figure 7D)** and liver **(Figure 7E)** of AKG treated mice. This was accompanied by an improved lung pathology upon AKG supplementation **(Figure 7F)** and a concomitant decrease in granuloma number **(Figure 7G)**. Finally, we observe a reduction in the levels of lipid peroxide adducts (MDA) in lungs of mice given AKG, suggesting a role for AKG in mitigating LPO induced during Mtb infection and hence, limiting dissemination **(Figure 7H)**. This sheds light on the therapeutic potential of AKG against TB pathogenesis.

**Figure 7:**
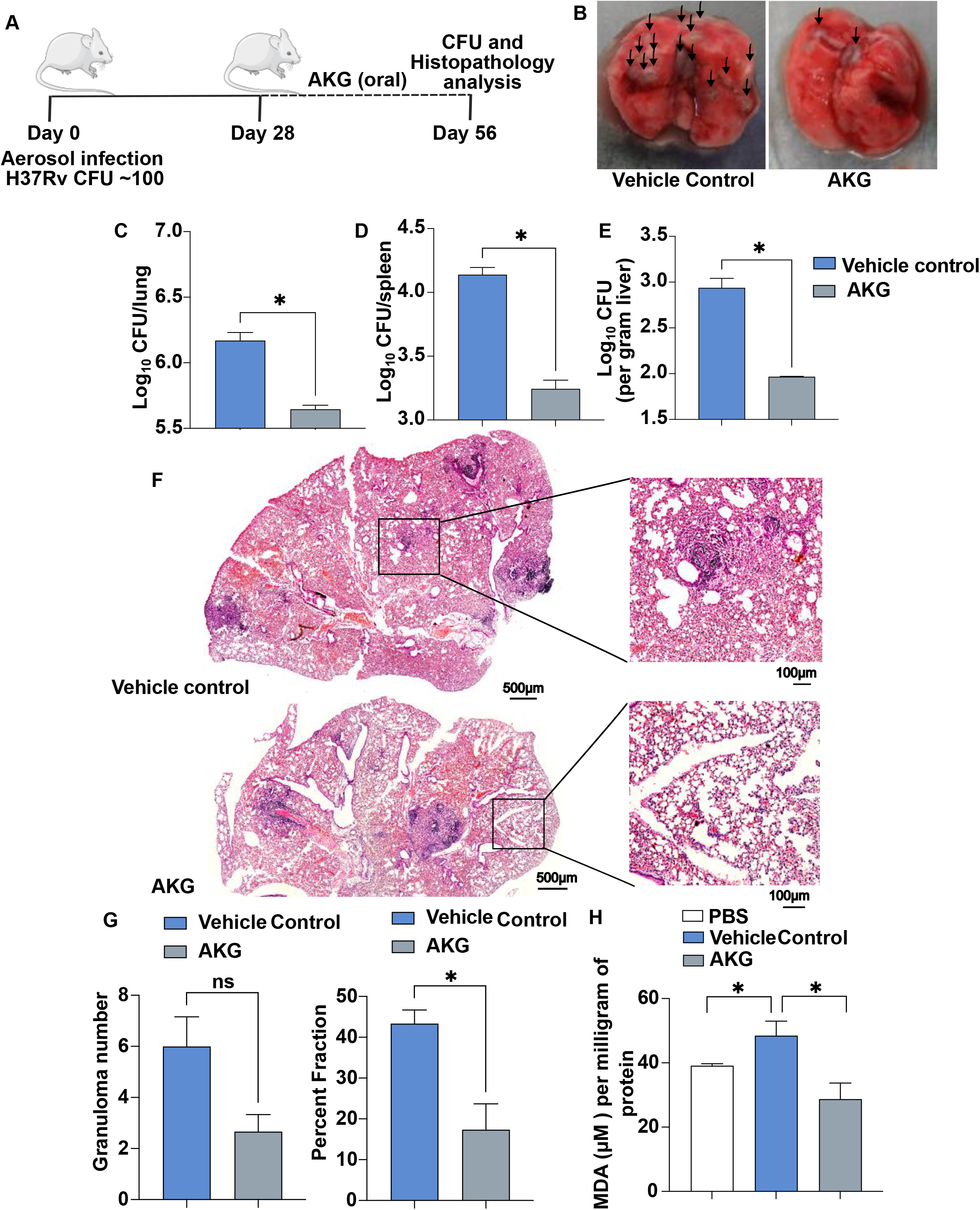
Dietary AKG as a supplement against TB pathogenesis. **(A-H)** Aerosol Mtb H37Rv infection (∼100 CFU) was given to mice. 4 weeks post infection, 1% alpha-ketoglutarate (AKG) was supplemented in drinking water (ad libitum) for 4 weeks. **(A)** schematic of the *in vivo* infection model used .**(B)** Gross lung pathology of murine lungs from infected mice with and without AKG supplementation. **(C-E)** CFU was enumerated from the **(C)** lungs, **(D)** spleen and **(E)** liver of AKG treated and untreated Mtb-infected mice. **(F)** H & E staining; representative image **(G)** Granuloma number (left), Percent fraction of lungs overed by granuloma (Right) **(H)** Levels of malondialdehyde (Lipid peroxidation byproduct) in Mtb H37Rv infected mice lungs given AKG supplementation. ns, non-significant *, p<0.05 Student’s t-test: C, D, E, G and H. N=3.

## DISCUSSION

Tuberculosis accounted for 1.23 million deaths in 2024 alone and as per WHO report 2025, infections with drug resistant strains of TB are on the rise^56^. The causative agent, *Mycobacterium tuberculosis*, a highly adaptive pathogen, continues to manipulate host cellular pathways to evade immune responses^57^. Mtb mediated hijacking of host transcriptional responses via regulating epigenetic landscape and rewiring of host metabolome have garnered much attention in the recent times^4^. In our study, we identify LSD1 as a key metabolic regulator during Mtb infection.

Depending on the site it acts on, LSD1 demethylase activity can lead to gene repression or activation and has been implicated in regulating various cellular processes^14^. LSD1 alters cellular metabolic state to support cancer growth and metastasis^15,31^ and hence, LSD1 inhibition as a cancer therapy is being evaluated with various inhibitors under clinical trials^19,20,58,59^. We report a pro-pathogen role of LSD1 during Mtb pathogenesis, as LSD1 was induced upon Mtb infection and its knockdown led to decreased mycobacterial survival. This is in line with previous research wherein LSD1 has been implicated in aiding replication of SARS-CoV2 ^60^ and Hepatitis C virus^61^, modulating host lipid metabolism to support *Cryptococcus neoformans* infection^16^. Our question was to discern LSD1 mediated changes induced by Mtb to support its intracellular survival.

Rewiring of host metabolome as a pathogen strategy to support infection and persistence has been implicated in various infections^62,63^. In case of Mtb infection, research has alluded to diverse metabolic adaptations including elevated glycolysis^64^, balanced OXPHOS-Glycolysis^65^ as well as suppression of both the pathways^66,67^ and recent research has shed light on the presence of metabolic heterogeneity among Mtb infected macrophages harboring different Mtb populations governing infection outcomes^35^. LSD1 acts as a regulator of metabolic shift, supporting glycolysis in hepatocellular carcinoma cells with its inhibition leading to increased OXPHOS and mitochondrial respiration^68^ and is known to promote tumor progression via facilitating adaptation of cancer cells to different microenvironments. With the aim to understand the metabolic state of Mtb infected macrophages upon LSD1 inhibition that limits mycobacterial survival, we carried out targeted metabolomics. Although our results did not indicate an overall change in host energy metabolism upon LSD1 knockdown during Mtb infection, we observed increase in the levels of AKG upon LSD1 knockdown in Mtb infected macrophages. This was also reflected in mice model of TB infection wherein LSD1 inhibitor treatment led to increased levels of AKG in lungs concomitant with improved lung pathology. This made us shift our focus to understanding LSD1 driven regulation of AKG and implications for the TCA metabolite in TB pathogenesis.

A key metabolite, AKG is generated either during TCA cycle from isocitrate by oxidative decarboxylation or via breakdown of the amino acid, glutamine (glutaminolysis). Immunological roles of AKG are diverse, generally associated with resolution of inflammation and infection^69^. Increased expression of glutamine utilization genes (*Gls, Glud1, Slc1a5, Slc1a3*) and increase in the levels of ^13^C_6_ derived AKG (labelled glucose) and ^12^C_6_ derived AKG (from glutamine) in our metabolic flux experiment upon LSD1 inhibition during Mtb infection indicate involvement of both pathways in AKG generation. Furthermore, inhibiting glutamine breakdown supported Mtb survival within the macrophages, pointing towards a crucial role for AKG in restricting mycobacterial survival.

AKG is implicated in redox metabolism, amino acid synthesis, regulating immune responses, oxidative stress, nitrogen metabolism as well as influencing host epigenetics and is generally associated with reducing chronic inflammation and benefitting overall human health^55^. AKG is a known antioxidant acting via various mechanisms: reducing the levels of Lactate dehydrogenase (LDH) which improves activity of antioxidative enzymes (eg. SOD, CAT, GSH) known to prevent lipid peroxidation; as an antagonist of cyanide and reducing cyanide induced oxidative stress; acts as a direct ROS scavenger during formation of succinate via quenching of hydrogen peroxide and hence mitigates H_2_O_2_ induced stress; increasing glutathione (GSH) levels^24^. There is contrasting evidence regarding the role of oxidative stress during Mtb pathogenesis. Several studies underscore ROS generation to be crucial for host defense and detail the various strategies employed by Mtb to mitigate oxidative stress and ensure its survival^70–72^. On the other hand, it is becoming increasingly clear that sustained ROS production and oxidative stress can drive lipid peroxidation, a process accelerated by increased labile iron pool and reduced activity of GPX4, resulting in accumulation of lipid peroxides and MDA adducts, ultimately leading to ferroptosis, an iron mediated cell death pathway^73^. Emerging evidence suggests that Mtb actively tries to increase iron levels in the cells to orchestrate ferroptosis which has been widely implicated to support its pathogenesis and dissemination^25–28^.

To test our hypothesis if increased AKG levels could counteract Mtb induced host lipid peroxidation, we carried out AKG supplementation during Mtb infection. Indeed, our results show that AKG supplemented macrophages showed reduced oxidative stress, decreased levels of MDA, increased GPX4 transcription and hence, consequent decrease in host LPO. Consistent with this, a direct role of AKG in inhibiting host ferroptosis and reducing MDA levels has previously been reported^29,30,74^. AKG supplementation also led to decreased intracellular survival of Mtb and reduced macrophage cell death, highlighting a protective role of AKG during Mtb infection. Furthermore, blocking glutamine breakdown reversed the effects of LSD1 inhibition mediated reduction in LPO and Mtb CFU, suggesting that LSD1 acts upstream of glutamine breakdown and regulates AKG levels to support Mtb infection.

Notably, AKG supplementation as a therapy has been evaluated against various infectious and metabolic diseases with promising results^55^. AKG has been shown to regulate aging^53^, alleviate ammonia induced lung injury^43^ and osteoarthiritis^30^, suppress the progression of non-alcoholic fatty liver disease ^40^, reduce thrombosis and inflammation associated with diabetes^52^, exert anti-tumor effects^48^ as well as mitigate lung inflammation during SARS CoV-2 infection^42^. In line with these reports, supplementation of AKG in Mtb infected mice resulted in marked decrease in lung bacillary burden, improved lung pathology as well as limited dissemination to extra pulmonary organs such as spleen and liver. Therefore, our results highlight AKG supplementation as a potential host-directed adjunct therapy against Tuberculosis.

Taken together, our study uncovers an epigenetic-metabolic axis exploited by Mtb via inducing LSD1 to ensure its survival within the host. LSD1 inhibition leads to increased levels of the TCA metabolite, AKG, via promoting glutamine breakdown. This increase in AKG levels suppresses host oxidative stress and lipid peroxidation via regulation of GPX4 and results in improved lung pathology, enhanced mycobacterial clearance and decreased Mtb dissemination **(Figure 8)**. Hence, our findings highlight the potential of AKG supplementation as a promising therapeutic strategy to control Tuberculosis.

**Figure 8:**
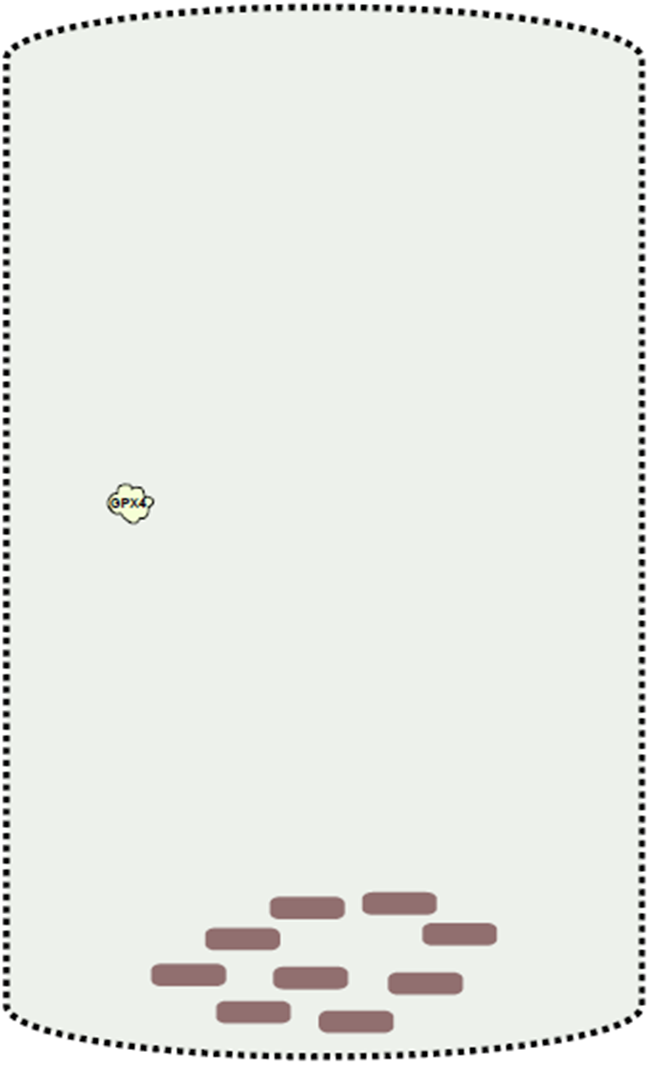
LSD1 modulates host AKG levels to support mycobacterial pathogenesis by regulating Lipid Peroxidation. Infection with *Mycobacterium tuberculosis* (Mtb) leads to increase in the levels of Lysine Specific Demethylase 1 (LSD1) that supports mycobacterial survival. Knocking down LSD1 increases expression of *Glutaminase (Gls)* implicated in glutamine (Gln) breakdown and results in an increase in the levels of the TCA metabolite, Alpha-ketoglutarate (AKG). AKG, being an antioxidant, suppresses Mtb induced Reactive Oxygen Species (ROS) generation and Lipid Peroxidation (LPO) via regulating Glutathione peroxidase 4 (GPX4), thus decreasing the accumulation of Malondialdehyde (MDA) adducts, a by-product of LPO. Consequently, LSD1 knockdown reduces intracellular survival of Mtb via attenuating LPO, a process known to facilitate Mtb pathogenesis and dissemination.

## MATERIAL AND METHODS

### Ethics statement

All mice experiments have been carried out after obtaining necessary approvals from Institutional Ethics Committee for Animal Experimentation (CAF/ethics/994/2023) and infection experiments using virulent *Mycobacterium tuberculosis* H37Rv from the Institutional Biosafety Committee (IBSC), Ref: IBSC/IlSc/KNB /05/2023-24. Protocols followed were in compliance with the guidelines set by the CCSEA (Committee for Control and Supervision of Experiments on Animals), Government of India.

### Cells and mice

Mice peritoneal macrophages used for *ex-vivo* experimentation were isolated four days after thioglycolate (HIMEDIA) injection from mice peritoneal cavity and maintained in 10% of Foetal Bovine Serum (FBS, Gibco) containing Dulbecco’s Minimal Eagle Medium (DMEM, Gibco) with 5% CO2 at 37°C. Metabolic experiments were carried out in mice Bone Marrow Derived Macrophages (BMDMs) by single-cell suspension preparation by flushing bone marrow from femurs and tibias in ice-cold 10% FBS-DMEM. Cells cultured in DMEM medium were supplemented with murine M-CSF at concentration of 25 ng/mL (Peprotech, Gibco) for differentiation. At 3^rd^ and 5^th^ day, media was replenished and experiments were carried out in differentiated cells post 7 days.

### Bacteria

Mtb H37Rv and tdTomato Mtb H37Rv virulent mycobacteria strains were received as a kind research gift from Prof. Amit Singh, IISc. Mtb cultures were grown in Middlebrook 7H9 medium (Difco, USA) supplemented with 10% OADC (oleic acid, albumin, dextrose, catalase) and hygromycin for tdTomato Mtb H37Rv to mid-log phase. Post 1X PBS washes, single-cell suspensions were obtained by passing through 23-, 28- and 30-gauge needle 10 times each and used at a multiplicity of infection 10 unless otherwise stated. All studies with virulent mycobacterial strains were conducted at the biosafety level 3 (BSL-3) facility at CIDR, IISc.

### Antibodies and Reagents

Chemicals and reagents: Sigma-Aldrich, Promega and HiMedia. Tissue culture plastic ware: Nunc Cell Culture, Thermo Fisher Scientific and Tarsons India Pvt. Ltd. siRNAs against *Kdm1a* and *Gls* were obtained from Eurogentec (sequence provided in Supplemental Table 1), Lipofectamine 3000 (L3000015): Thermo Fisher Scientific. Rabbit Anti-LSD1 (2139; RRID:AB_2070135): Cell Signaling Technology; DAPI (4′,6-Diamidino-2-phenylindole dihydrochloride): Thermo Fisher Scientific (D1306); anti-MAP1LC3B (L7543) : Sigma Aldrich; anti-β-ACTIN HRP-tagged (A3854), anti-rabbit IgG/anti-mouse IgG conjugated with HRP: Jackson, USA.

### Pharmacological reagents

Following 1h pre-treatment with LSD1 inhibitor (Sigma M74253)-3μM, Alpha-ketoglutarate (Sigma K1128) -2mM., cells were used for infection studies. PBS was utilised as vehicle control.

### Transient transfections

Lipofectamine-3000 reagent (Invitrogen) and Opti-MEM (Gibco) with 100nM of either non-targeting or specific siRNAs was utilized for carrying out transfections.

### Real-time qRT-PCR

TRI Reagent (Sigma) was used for macrophage RNA isolation and 1μg was utilised for cDNA synthesis using synthesis kit. Using SYBR Green PCR mix (Takara), Quantitative real time PCR for analysis of target gene expression with *Gapdh* as an internal control was set up. Fold change was calculated via comparative ΔΔCt method. Supplemental Table 2 provides sequences of primers used.

### Immunoblotting

Post infection/ treatment, cells lysed in RIPA buffer [0.25% sodium-deoxycholate, 150 mM NaCl, 50 mM Tris-HCl pH 7.4, 1% NP-40, 1 mM EDTA, 1 mM PMSF, and protease inhibitor cocktail (1mM): Pepstatin, Leupeptin, NaF, Aprotinin and Na3VO4 (Sigma). Protein concentration was estimated by Bradford’s assay (Sigma) and SDS-PAGE was carried out in 5% stacking and 12% resolving gel. The proteins were then transferred onto PVDF (polyvinylidene difluoride) membranes (Millipore). Blocking of the membrane was done for 1h in milk powder (5%) (HIMEDIA) prepared in 0.1% Tween 20 with 20mM Tris-HCl (pH 7.4) and 137mM NaCl (TBST) (Sigma) and primary antibody (5% BSA, MP Bio) was probed overnight at 4°C. TBST was used to wash the blots followed by secondary Ab incubation (HRP conjugated goat anti-rabbit IgG). Enhanced chemiluminescence protocol (BIO-RAD) utilized to develop the blots and β-ACTIN was loaded as control.

### Metabolite extraction and LC-MS/MS analysis

Post 24h infection with Mtb, BMDMs with and without siRNA mediated knockdown of LSD1 were processed for mass spectrometry via extraction of polar metabolites as per an established protocol^37^. Briefly, the cells were harvested in 1X PBS and cell pellet was dissolved in 75% (v/v) ethanol (600μL) and internal standard for derivatized metabolites (TCA cycle) [2 nmol of D4-succinic acid (Cambridge Isotopes, catalog# DLM-2307). Following incubation of 3 min at 80°C and 5 min in ice, samples centrifuged at 20,000*g* for 10 min at 4 °C. Polar metabolites containing supernatant transferred to a new tube, vacuum dried, stored for further LC-MS/MS analysis at −80°C.

For processing,150μl of water and 75μL of 1M N-(3-dimethylaminopropyl)-N′-ethyl carbodiimide in 13.5 mM pyridine buffer at pH 5.0 (Sigma, catalog# E7750) and 150μL of 0.5M *O-*benzylhydroxylamine (Sigma, catalog# B22984), in 13.5 mM pyridine buffer at pH 5.0, was added and incubation of 1h at 25°C. Next, 350μl of ethyl acetate was added, shaking for 10min and centrifugation at 3,000*g* for 5 min at 4°C. Ethyl acetate extraction carried out thrice and top layers collected from each round pooled together, vacuum dried, and stored for further LC-MS/MS analysis at −80°C. Similar metabolite extraction was carried out for *in vivo* samples wherein lung and liver isolated from the mice were flash frozen and stored in -80°C. 100 mg of each organ was homogenized in PBS followed by ethanol extraction.

For the metabolic flux assay, BMDMs were differentiated and maintained in normal DMEM (4mM glutamine, 25mM glucose) to reach steady state and siRNA treatment was carried out followed by starvation phase where cells were maintained in DMEM with no glucose for 2h. Media was replaced with no glucose DMEM + 25mM 13C6 Glucose (+10% dialyzed FBS) and infected with H37Rv at MOI 1:10 and cells harvested at mentioned time points. Post PBS wash, cells were directly quenched and metabolite extraction carried out as detailed above. Agilent 6545 Q-TOF mass spectrometer fitted with an Agilent 1290 Infinity II UHPLC system and AutoMS/MS acquisition method was utilized for LC-MS runs following published protocols^37,75^. Re-solubilization of dried metabolites was done in appropriate solvents. For non-derivatized metabolites, a mixture of H_2_O:Acetonitrile (19:1) and 5 mM Ammonium Acetate was used. The derivatized metabolites were solubilized in 50 µL of MeOH:H_2_O (1:1). 10μL of the resuspended samples injected onto a Phenomenex Synergi Fusion-RP Column of specifications 150 mm x 4.6 mm, 4 μm, 80 Å (catalog # 00F-4424-E0) fitted with a Phenomenex guard column of specification 3.2 mm X 8.0 mm (catalog # KJ0-4282).

For metabolite data analysis, all the peaks manually validated based on fragments obtained, if any, and relative retention times. Agilent MassHunter Qualitative Analysis 10.0 software was used. A mass accuracy of 15 ppm was maintained for all detected species and area under the curve for different metabolites was measured normalized to internal standard (D4-succinic acid) levels and cell number or weight of tissue.

### Flow cytometry analysis

Following infection and treatment, cells gently scraped in 1X PBS were stained with a) antibody for LSD1 (CST) and Alexa488-conjugated antibody at room temperature for 30min b) CellROX Deep Red Reagent at room temperature for 1h c) BODIPY C11 dye for lipid peroxidation. Flow cytometry was carried out in BC cytoFLEX followed by analysis in FlowJo software.

### BODIPY Staining

Post infection/treatment, 3.7% formaldehyde fixation of the cells for 30 min and staining with 10 μg/mL BODIPY 493/503 (Invitrogen, D3922) and BODIPY 665/676 (Invitrogen, B3932) for 30 min, followed by nuclei staining with DAPI and glycerol mounting. Confocal microscopy with Zeiss LSM 880 confocal laser scanning microscope (Carl Zeiss AG, Germany) using a plan-Apochromat 63X/1.4 Oil DIC objective (Carl Zeiss AG, Germany) was carried out. For quantification, measurement of the area-integrated intensity was carried out in ImageJ using the free hand selection tool. CTCF, Corrected Total Cell Fluorescence: (area of selected cell X Mean florescence intensity) – (area of selected cell X Background mean florescence intensity). Fluorescence of an area without cells in the field was used to obtain the background intensity. CTCF (BODIPY 665/676)/ CTCF (BODIPY 493/503) gave us the proportion of unoxidized and neutral lipids.

### PI staining

Macrophage cell death was assessed using Propidium Iodide (PI) staining. Following infection/treatment and washes with 1X PBS, PI staining (5µg/ml) for 5min at room temperature. Next, cell fixation with 3.7% formaldehyde for 30min, nuclei staining with DAPI and glycerol mounting (Sigma). Next, confocal microscopy was done as described above. Counting the DAPI positive nuclei gave number of total cells in a field with PI positive nuclei counted as dead macrophages. Dead cell percentage per field was calculated as (PI positive cells/ DAPI positive cells)*100. For each condition, 20-30 randomly selected fields with minimum of 50 cells per field were analysed.

### Immunofluorescence

3.7% formaldehyde was used to fix cells post infection and treatment followed by blocking for 1h using 0.02% saponin (Sigma) in 2% BSA and incubated with LSD1 antibody overnight at 4°C and 30 min with Alexa488-conjugated secondary antibody at room temperature with counter staining of nucleus with DAPI, glycerol mounting and processed for imaging and analysis.

### TBARS assay

TBARS assay kit (Cayman Chemicals, 703002) was utilized and the kit protocol was followed. To homogenized lung tissues or cells lysed in RIPA buffer, colour reagent and SDS solution was added followed by incubation for 1h at 95°C. After cooling of the samples to room temperature, centrifugation was carried out. Microplate reader (SpectraMax M3) was used to measure absorbance at 532 nm. For determination of the amount of MDA (μM) in the samples, standard curve was drawn.

### Intracellular CFU of Mtb

Peritoneal macrophages post treatment with the mentioned inhibitor/siRNA were given Mtb H37Rv infection at MOI 5. Post 4h internalization, cells were maintained in 0.2 mg/ml amikacin containing media for 2h for killing of extracellular mycobacteria. Following this, 7H9 media was used to wash the cells and one set was taken as 0 h and other replicates maintained in medium without antibiotics for mentioned time. Lysis with 0.06% SDS in 7H9 media was carried out and in oleic acid, albumin, dextrose, catalase (OADC) supplemented Middlebrook 7H11 agar, plated in appropriate dilutions. After 21 days, intracellular mycobacteria was enumerated by counting total colony forming units (CFUs).

### *In vivo* infection study

Infection with Mtb H37Rv at 100 CFU/animal was given to BALB/c mice in a Madison chamber instrument inside a securely commissioned BSL3 facility and maintained for 28 days. Post this, depending on the experiment design, mice were either treated with LSD1 inhibitor (50mg/Kg, Sigma) intra-peritoneally, Isoniazid (10mg/kg) in water or 1% AKG was supplemented in water for 28 days. In each case, mice were sacrificed on the 56^th^ day and CFU was enumerated from spleen, liver and left lobe of the lung via homogenization in sterile PBS and serial dilution for plating on 7H11 agar containing OADC. Lower right lobes were flash frozen and stored in -80 for MDA assay and metabolite extraction and upper right lobes of the lungs were embedded in paraffin post formalin fixation for histopathology analysis.

### H/E staining

Paraffin-embedded lung tissue samples were sliced into 5μm thick sections using Leica RM2245 microtome. H and E staining was done in deparaffinized and rehydrated sections. This was followed by dehydration and tissue samples were mounted on coverslip using DPX. The samples were then sent for a blinded pathologist analysis for granuloma scoring and quantification of lung fraction involved in pathology.

### Statistical analysis

All tests used in respective experiments have been specified in the figure legends along with p-values. In general, one-way ANOVA followed by Tukey’s multiple-comparisons was used for assessing multiple samples and Student’s t-test for two samples. P values < 0.05 were defined as significant with data in the graphs plotted as the mean ± SEM and representative of 3 independent experiments. Graphing and analysis was done using GraphPad Prism 10.0 software (GraphPad Software, USA).

## ACKNOWLEDGEMENT

We would like to thank the Central Animal Facility, IISc for providing mice for experimentation and BSL3 facility at CIDR, IISc for infection studies with Mtb H37Rv. We are grateful to Prof. Amit Singh for insightful discussions and suggestions. We acknowledge the help of Prof. K.N. Balaji’s research group, specifically Smriti Sundar for her valuable inputs during manuscript preparation. The schematics have been made using MS-PowerPoint and InkScape with freely available images.

## AUTHOR CONTRIBUTIONS

Conceptualization, A.S., and K.N.B.; methodology, A.S., A.A. and K.N.B.; investigation, A.S., A.A., A.C., R.S.R. and S.S.K; writing—original draft, A.S., and K.N.B.; writing—review and editing, A.S., A.A. and K.N.B.; funding acquisition, K.N.B.; resources, K.N.B.; supervision, K.N.B.

## REFERENCES

1. Bierne, H., Hamon, M. & Cossart, P. Epigenetics and bacterial infections. Cold Spring Harb. Perspect. Med. 2, (2012).

2. Fol, M., Włodarczyk, M. & Druszczyńska, M. Host Epigenetics in Intracellular Pathogen Infections. International Journal of Molecular Sciences 2020, Vol. 21, Page 4573 21, 4573 (2020).

3. Kathirvel, M. & Mahadevan, S. The role of epigenetics in tuberculosis infection. Epigenomics 8, 537–549 (2016).

4. Khadela, A. et al. Epigenetics in Tuberculosis: Immunomodulation of Host Immune Response. Vaccines (Basel). 10, (2022).

5. Qu, L. et al. Histone demethylases in the regulation of immunity and inflammation. Cell Death Discovery 2023 9:1 9, 188- (2023).

6. Chen, Y. C. et al. Histone H3K14 hypoacetylation and H3K27 hypermethylation along with HDAC1 up-regulation and KDM6B down-regulation are associated with active pulmonary tuberculosis disease. Am. J. Transl. Res. 9, 1943 (2017).

7. Holla, S. et al. MUSASHI-Mediated Expression of JMJD3, a H3K27me3 Demethylase, Is Involved in Foamy Macrophage Generation during Mycobacterial Infection. PLoS Pathog. 12, e1005814 (2016).

8. Kim, D., Kim, K. Il & Baek, S. H. Roles of lysine-specific demethylase 1 (LSD1) in homeostasis and diseases. J. Biomed. Sci. 28, 41 (2021).

9. Yang, Y. T., Wang, X., Zhang, Y. Y. & Yuan, W. J. The histone demethylase LSD1 promotes renal inflammation by mediating TLR4 signaling in hepatitis B virus-associated glomerulonephritis. Cell Death & Disease 2019 10:4 10, 278- (2019).

10. Li, Y. et al. Lipidomic profiling reveals lipid regulation by a novel LSD1 inhibitor treatment. Oncol. Rep. 46, (2021).

11. Musri, M. M. et al. Histone demethylase LSD1 regulates adipogenesis. Journal of Biological Chemistry 285, 30034–30041 (2010).

12. Sakamoto, A. et al. Lysine demethylase LSD1 coordinates glycolytic and mitochondrial metabolism in hepatocellular carcinoma cells. Cancer Res. 75, 1445–1456 (2015).

13. Wang, Z. et al. Reprogramming of glutamine metabolism by epigenetic modifier LSD1 regulates chemotherapy resistance in PDAC. Pancreatology 21, S67 (2021).

14. Maiques-Diaz, A. & Somervaille, T. C. P. LSD1: Biologic roles and therapeutic targeting. Epigenomics vol. 8 1103–1116 Preprint at 10.2217/epi-2016-0009 (2016).

15. Fernandez-Zapico, M. E., Kozub, M. M., Carr, R. M. & Lomberk, G. L. LSD1, a double-edged sword, confers dynamic chromatin regulation but commonly promotes aberrant cell growth. F1000Research vol. 6 Preprint at 10.12688/f1000research.12169.1 (2017).

16. Lohia, G. K., Shah, A. & Balaji, K. N. Histone demethylase LSD1 regulates lipid homeostasis during Cryptococcus neoformans infection. iScience 28, 113405 (2025).

17. Shan, J. et al. Histone demethylase LSD1 restricts influenza A virus infection by erasing IFITM3-K88 monomethylation. PLoS Pathog. 13, (2017).

18. Shan, J. et al. Histone demethylase LSD1 restricts influenza A virus infection by erasing IFITM3-K88 monomethylation. PLoS Pathog. 13, (2017).

19. Wass, M. et al. A proof of concept phase I/II pilot trial of LSD1 inhibition by tranylcypromine combined with ATRA in refractory/relapsed AML patients not eligible for intensive therapy. Leukemia 2020 35:3 35, 701–711 (2020).

20. Noce, B., Di Bello, E., Fioravanti, R. & Mai, A. LSD1 inhibitors for cancer treatment: Focus on multi-target agents and compounds in clinical trials. Front. Pharmacol. 14, (2023).

21. Zwergel, C., Stazi, G., Mai, A. & Valente, S. Trends of LSD1 inhibitors in viral infections. Future Medicinal Chemistry vol. 10 1133–1135 Preprint at 10.4155/fmc-2018-0065 (2018).

22. Wu, N. et al. Alpha-Ketoglutarate: Physiological Functions and Applications. Biomol. Ther. (Seoul). 24, 1 (2016).

23. Liu, S., Yang, J. & Wu, Z. The Regulatory Role of α-Ketoglutarate Metabolism in Macrophages. Mediators Inflamm. 2021, (2021).

24. Liu, S., He, L. & Yao, K. The Antioxidative Function of Alpha-Ketoglutarate and Its Applications. Biomed Res. Int. 2018, 3408467 (2018).

25. Qiang, L. et al. A mycobacterial effector promotes ferroptosis-dependent pathogenicity and dissemination. Nature Communications 2023 14:1 14, 1430- (2023).

26. Amaral, E. P. et al. A major role for ferroptosis in Mycobacterium tuberculosis–induced cell death and tissue necrosis. Journal of Experimental Medicine 216, 556–570 (2019).

27. Satish, B. A., Sundar, S., Rajmani, R. S. & Balaji, K. N. Activin A mediated KAT8 expression induces ferroptosis during Mycobacterium tuberculosis infection. J. Infect. Dis. https://doi.org/10.1093/INFDIS/JIAF482 (2025) doi:10.1093/INFDIS/JIAF482.

28. Sundar, S., Rajmani, R. S. & Balaji, K. N. Infection with Mycobacterium tuberculosis orchestrates the PRMT5-dependent methylation of NCOA4 to govern host ferroptosis. bioRxiv 2025.10.23.684158 (2025) doi:10.1101/2025.10.23.684158.

29. Yu, H. et al. α-Ketoglutarate improves cardiac insufficiency through NAD+-SIRT1 signaling-mediated mitophagy and ferroptosis in pressure overload-induced mice. Molecular Medicine 30, 15 (2024).

30. He, R. et al. α-Ketoglutarate alleviates osteoarthritis by inhibiting ferroptosis via the ETV4/SLC7A11/GPX4 signaling pathway. Cell. Mol. Biol. Lett. 29, (2024).

31. Ding, J. et al. LSD1-mediated epigenetic modification contributes to proliferation and metastasis of colon cancer. Br. J. Cancer 109, 994–1003 (2013).

32. Sobczak, M. et al. Lsd1 facilitates pro-inflammatory polarization of macrophages by repressing catalase. Cells 10, (2021).

33. Kim, D. et al. PKCα-LSD1-NF-κB-Signaling Cascade Is Crucial for Epigenetic Control of the Inflammatory Response. Mol. Cell 69, 398-411.e6 (2018).

34. Cao, Y. et al. Anterograde regulation of mitochondrial genes and FGF21 signaling by hepatic LSD1. JCI Insight 6, (2021).

35. Yadav, V. et al. Bioenergetic reprogramming of macrophages reduces drug tolerance in Mycobacterium tuberculosis. Nature Communications 2025 16:1 16, 9370- (2025).

36. Walvekar, A., Rashida, Z., Maddali, H. & Laxman, S. A versatile LC-MS/MS approach for comprehensive, quantitative analysis of central metabolic pathways. Wellcome Open Res. 3, (2018).

37. Rajendran, A., Soory, A., Khandelwal, N., Ratnaparkhi, G. & Kamat, S. S. A multi-omics analysis reveals that the lysine deacetylase ABHD14B influences glucose metabolism in mammals. Journal of Biological Chemistry 298, 102128 (2022).

38. Wu, N. et al. Alpha-Ketoglutarate: Physiological Functions and Applications. Biomol. Ther. (Seoul). 24, 1 (2016).

39. Wu, G. et al. Metabolism-driven posttranslational modifications and immune regulation: Emerging targets for immunotherapy. Science Advances 11, 1–19 (2025).

40. Nagaoka, K. et al. The metabolite, alpha-ketoglutarate inhibits non-alcoholic fatty liver disease progression by targeting lipid metabolism. Liver Res. 4, 94–100 (2020).

41. Cheng, D. et al. α-Ketoglutarate prevents hyperlipidemia-induced fatty liver mitochondrial dysfunction and oxidative stress by activating the AMPK-pgc-1α/Nrf2 pathway. Redox Biol. 74, (2024).

42. Agarwal, S. et al. Dietary alpha-ketoglutarate inhibits SARS CoV-2 infection and rescues inflamed lungs to restore O2 saturation by inhibiting pAkt. Clin. Transl. Med. 12, e1041 (2022).

43. Ali, R., Mittal, G., Sultana, S. & Bhatnagar, A. Ameliorative potential of alpha-ketoglutaric acid (AKG) on acute lung injuries induced by ammonia inhalation in rats. Exp. Lung Res. 38, 435–444 (2012).

44. Liang, G. et al. α-Ketoglutarate plays an inflammatory inhibitory role by regulating scavenger receptor class a expression through N6-methyladenine methylation during sepsis. Eur. J. Immunol. 54, 2350655 (2024).

45. Liu, S., He, L. & Yao, K. The Antioxidative Function of Alpha-Ketoglutarate and Its Applications. Biomed Res. Int. 2018, 3408467 (2018).

46. He, L. et al. Prevention of Oxidative Stress by α-Ketoglutarate via Activation of CAR Signaling and Modulation of the Expression of Key Antioxidant-Associated Targets in Vivo and in Vitro. J. Agric. Food Chem. 66, 11273–11283 (2018).

47. Amaral, E. P. et al. A major role for ferroptosis in Mycobacterium tuberculosis-induced cell death and tissue necrosis. J. Exp. Med. 216, 556–570 (2019).

48. Greilberger, J., Herwig, R., Greilberger, M., Stiegler, P. & Wintersteiger, R. Alpha-Ketoglutarate and 5-HMF: A Potential Anti-Tumoral Combination against Leukemia Cells. Antioxidants (Basel) 10, (2021).

49. Tian, Q. et al. Dietary Alpha-Ketoglutarate Promotes Epithelial Metabolic Transition and Protects against DSS-Induced Colitis. Mol. Nutr. Food Res. 65, 2000936 (2021).

50. Long, F., Xiang, J., Tan, Y., Zhao, J. & Ma, C. The research progress of α-ketoglutarate in osteoarthritis. Biochem. Biophys. Rep. 44, 102270 (2025).

51. Shrimali, N. M. et al. α-Ketoglutarate Inhibits Thrombosis and Inflammation by Prolyl Hydroxylase-2 Mediated Inactivation of Phospho-Akt. EBioMedicine 73, (2021).

52. Agarwal, S., Ghosh, R., Verma, G., Khadgawat, R. & Guchhait, P. Alpha-ketoglutarate supplementation reduces inflammation and thrombosis in type 2 diabetes by suppressing leukocyte and platelet activation. Clin. Exp. Immunol. 214, 197 (2023).

53. Rhoads, T. W. & Anderson, R. M. Alpha-Ketoglutarate, the Metabolite that Regulates Aging in Mice. Cell Metab. 32, 323–325 (2020).

54. Naeini, S. H., Mavaddatiyan, L., Kalkhoran, Z. R., Taherkhani, S. & Talkhabi, M. Alpha-ketoglutarate as a potent regulator for lifespan and healthspan: Evidences and perspectives. Exp. Gerontol. 175, 112154 (2023).

55. Gyanwali, B. et al. Alpha-Ketoglutarate dietary supplementation to improve health in humans. Trends in Endocrinology & Metabolism 33, 136–146 (2022).

56. Global tuberculosis report 2025. https://www.who.int/publications/i/item/9789240116924.

57. Chai, Q., Wang, L., Liu, C. H. & Ge, B. New insights into the evasion of host innate immunity by Mycobacterium tuberculosis. Cellular & Molecular Immunology 2020 17:9 17, 901–913 (2020).

58. Sattler, M. & Salgia, R. LSD1-targeted therapy—a multi-purpose key to unlock immunotherapy in small cell lung cancer. Transl. Lung Cancer Res. 12, 1350–1354 (2023).

59. Karakaidos, P., Verigos, J. & Magklara, A. LSD1/KDM1A, a Gate-Keeper of Cancer Stemness and a Promising Therapeutic Target. Cancers (Basel). 11, (2019).

60. Tu, W. J. et al. Targeting novel LSD1-dependent ACE2 demethylation domains inhibits SARS-CoV-2 replication. Cell Discovery 2021 7:1 7, 1–25 (2021).

61. Papadopoulou, G. et al. The Epigenetic Controller Lysine-Specific Demethylase 1 (LSD1) Regulates the Outcome of Hepatitis C Viral Infection. Cells 12, 2568 (2023).

62. Escoll, P. & Buchrieser, C. Metabolic reprogramming of host cells upon bacterial infection: Why shift to a Warburg-like metabolism? FEBS J. 285, 2146–2160 (2018).

63. Arumugam, P. & Kielian, T. Metabolism Shapes Immune Responses to Staphylococcus aureus. J. Innate Immun. 16, 12 (2023).

64. Lachmandas, E. et al. Rewiring cellular metabolism via the AKT/mTOR pathway contributes to host defence against Mycobacterium tuberculosis in human and murine cells. Eur. J. Immunol. 46, 2574–2586 (2016).

65. Gleeson, L. E. et al. Cutting Edge: Mycobacterium tuberculosis Induces Aerobic Glycolysis in Human Alveolar Macrophages That Is Required for Control of Intracellular Bacillary Replication. J. Immunol. 196, 2444–2449 (2016).

66. Cumming, B. M., Addicott, K. W., Adamson, J. H. & Steyn, A. J. C. Mycobacterium tuberculosis induces decelerated bioenergetic metabolism in human macrophages. Elife 7, (2018).

67. Olson, G. S. et al. Type I interferon decreases macrophage energy metabolism during mycobacterial infection. Cell Rep. 35, (2021).

68. Sakamoto, A. et al. Lysine Demethylase LSD1 Coordinates Glycolytic and Mitochondrial Metabolism in Hepatocellular Carcinoma Cells. Cancer Res. 75, 1445–1456 (2015).

69. Liu, S., Yang, J. & Wu, Z. The Regulatory Role of α-Ketoglutarate Metabolism in Macrophages. Mediators Inflamm. 2021, 5577577 (2021).

70. Pacl, H. T., Reddy, V. P., Saini, V., Chinta, K. C. & Steyn, A. J. C. Host-pathogen redox dynamics modulate Mycobacterium tuberculosis pathogenesis. Pathog. Dis. 76, fty036 (2018).

71. Shastri, M. D. et al. Role of Oxidative Stress in the Pathology and Management of Human Tuberculosis. Oxid. Med. Cell. Longev. 2018, 7695364 (2018).

72. Borbora, S. M. et al. Mycobacterium tuberculosis Elevates SLIT2 Expression Within the Host and Contributes to Oxidative Stress Responses During Infection. J. Infect. Dis. https://doi.org/10.1093/INFDIS/JIAD126 (2023) doi:10.1093/INFDIS/JIAD126.

73. Xie, Y. et al. Ferroptosis: process and function. Cell Death & Differentiation 2016 23:3 23, 369–379 (2016).

74. Guo, S. et al. Dietary α-ketoglutarate supplementation improves hepatic and intestinal energy status and anti-oxidative capacity of Cherry Valley ducks. Animal Science Journal 88, 1753–1762 (2017).

75. Agrawal, S. et al. A metabolic hierarchy directs cell cycle transition and morphogenesis. bioRxiv 2025.12.20.695365 (2025) doi:10.64898/2025.12.20.695365.

