## Supplemental Information for "Lysine specific demethylase 1 (LSD1) regulates host alpha-ketoglutarate levels to modulate lipid peroxidation during *Mycobacterium tuberculosis* infection"

### SUPPLEMENTAL FIGURES

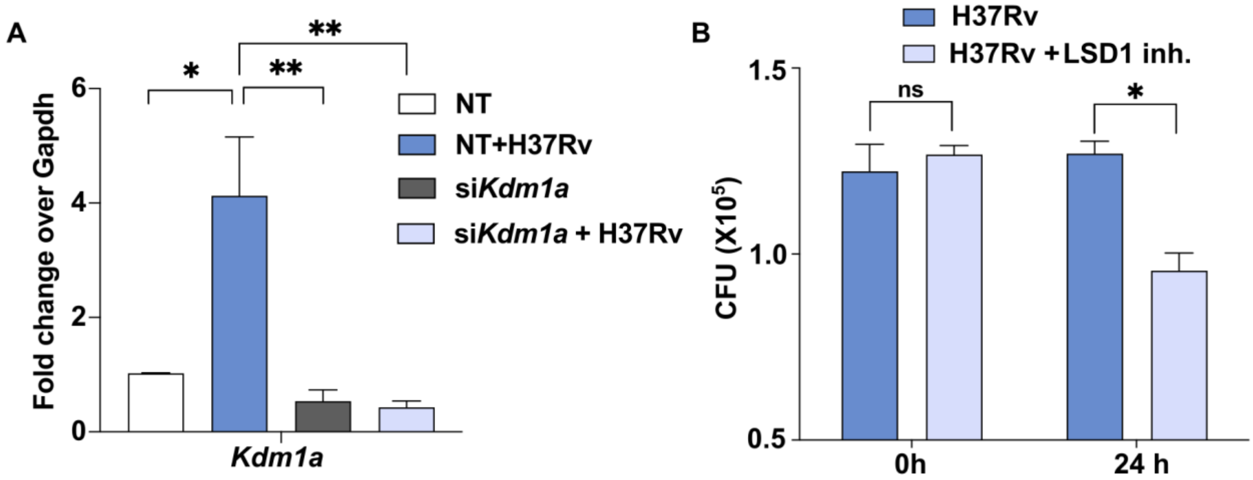


**Supplemental Figure 1: Decreased Mtb survival upon LSD1 inhibiton. (A)** BALB/c mouse peritoneal macrophages transiently transfected with siRNAs against *Kdm1a* and knockdown was validated at transcript level. **(B)** *In vitro* CFU was assessed 24 h post H37Rv infection (MOI 1:5) in BALB/c mouse peritoneal macrophages treated with LSD1 inhibitor (3 μM). *, p< 0.05 **, p< 0.01 One-Way ANOVA: A; ns, non-significant *, p< 0.05 Student's t-test: B

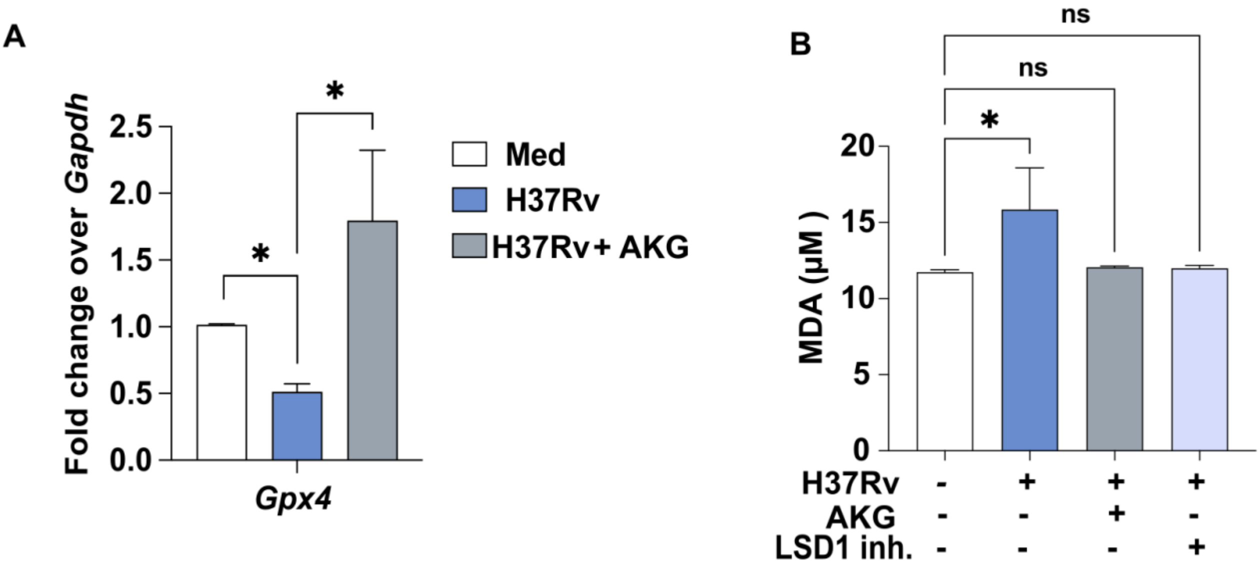


**Supplemental Figure 2: AKG regulates GPX4 mediated LPO during Mtb infection.** Mice macrophages infected with H37Rv for 24h with and without LSD1 inhibitor treatment or AKG supplementation were assessed for levels of (A) transcript levels of *Gpx4* and (B) malondialdehyde (Lipid peroxidation) by TBAR Assay; non-significant *, p< 0.05 One-Way ANOVA: A, B.

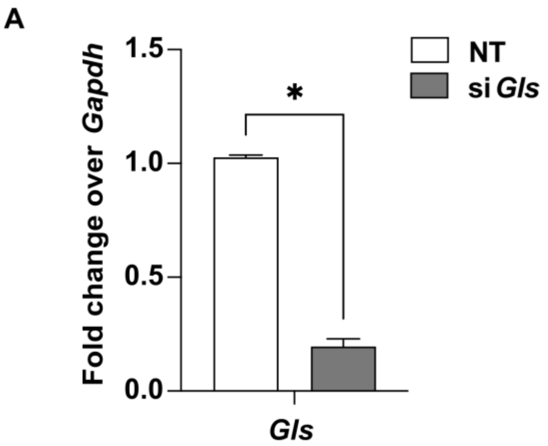


**Supplemental Figure 3: Validation of siRNA mediated knockdown of Glutaminase. (A)** BALB/c mouse peritoneal macrophages transiently transfected with siRNAs against *Gls* and knockdown was validated at transcript level. *, $p < 0.05$ Student's t-test.

### SUPPLEMENTAL TABLE 1: List of sequences for SiRNA mediated Knockdown

| ***Genes*** | **siRNA sequence** |
|---|---|
| *Kdm1a* | gacagacaaauacuugacu<br>gaagccacuucuuaccuua |
| *Gls* | tgttgtgacttctctaata<br>gcattcttgtggcatgtat |

### SUPPLEMENTAL TABLE 2: List of primers for mouse gene expression analyses

| ***Genes*** | **Forward primer (5'-3')** | **Reverse primer (5'-3')** |
|---|---|---|
| *Gapdh* | gagccaaacgggtcatcatct | Gaggggccatccacagtctt |
| *Kdm1a* | cttagagcgccatggtcttatc | tagcaactcgtccacctact |
| *Slc1a5* | gaacgaggtgtctctgaatc | cctcatccagtccatttctc |
| *Slc1a3* | tctctgggacagatccatac | taaacaagcacactccactc |
| *Slc7a5* | *gattctgcaggctatcttctc* | *gggacagtggattgtgttag* |
| *Gfpt2* | ctgaaccaactgccaagat | gcacagtagtggaaagtctg |
| *Gls* | ccgcagagtaagagaaatgg | gggagaaagagaacgactaaat |
| *Got1* | actcagggcaagactagaa | tggaggtagcgacgtaat |
| *Got2* | catagcctcctctctcatca | agtatctccttctgatcctacc |
| *Gpt2* | tcactccatctgtcctcttt | ctaagcaactccctgtctttag |
| *Glud1* | cgacttctttaccacctcttc | cacctaactggcttggatac |
| *Gpx4* | ggagccaggaagtaatcaag | cgcagccgttcttatcaa |
